# Hierarchical Bayesian modeling identifies key considerations in the development of quantitative loop-mediated isothermal amplification assays

**DOI:** 10.1101/2023.01.16.524143

**Authors:** Jacob R. Bradley, Diego Borges, Mafalda Cavaleiro, Michael B. Mayhew

## Abstract

**Motivation:** Loop-mediated isothermal amplification (LAMP) is a rapidly growing, fast, and cost-effective technique for detection of DNA/RNA in point-of-care biomedical applications. However, it remains unclear what factors affect LAMP’s quantitative resolution, and experimental optimization of primers presents a major bottleneck in assay design. A lack of model-based frameworks to characterize LAMP data and address these questions presents an unmet need for LAMP assay development.

**Results:** We present hierarchical Bayesian models of LAMP amplification based on Gompertz functions, and use these models to infer the effect of RNA variation and other factors on LAMP amplification curves derived from 80 blood samples of patients with suspected acute infection. Our analysis uncovers associations between LAMP assay resolution and characteristics such as primer sequence composition and thermodynamic properties. In addition to correlations between RNA input abundance and time shift of the the LAMP amplification curve, we also detect RNA-dependent assocations with amplification rate. We further investigate associations between primer/target properties and quantitative performance of the assay by generating a set of synthetic RNA samples with systematically varied primer sequences and applying our framework. We find evidence that the associations observed are driven by across-target rather than within-target variation, an important observation for study design. Our findings represent important first steps towards guided development of quantitative LAMP assays.

**Availability and Implementation:** Analysis and modeling code is available upon reasonable request.

## 1 Introduction

Loop-mediated isothermal amplification (LAMP) is a fast, sensitive, and precise method for detecting DNA or RNA (the latter case referred to as reverse-transcription (RT)-LAMP). Since its introduction (Notomi *et al*., 2000), LAMP has been used for pathogen detection (Mekata *et al*., 2009; Cao *et al*., 2017; Thiessen *et al*., 2018), sex identification (Hirayama *et al*., 2013; Almasi and Almasi, 2017; Centeno-Cuadros *et al*., 2018), and cancer monitoring (Li *et al*., 2016; Horiuchi *et al*., 2020; Kalofonou *et al*., 2020). Unlike comparable assays such as polymerase chain reaction (PCR), LAMP is isothermal and therefore does not require a thermocycler. This property makes LAMP amenable to point-of-care and field applications (Fu *et al*., 2011). As in PCR, LAMP specificity relies on primer-target complementarity. LAMP was originally conceived with two pairs of primers (F3/B3 and FIP/BIP; Figure 1), and further optional pairs of primers were introduced for loop hybridization (Nagamine *et al*., 2002) and amplicon enlargement (Gandelman *et al*., 2011). The need for coordinated design of multiple primers can create a major bottleneck in LAMP assay development.

While LAMP has primarily been used in qualitative/detection applications, advances in measurement technology (Zhang *et al*., 2014; Becherer *et al*., 2020), in particular automated fluorescence detection for producing LAMP amplification curves, have enabled quantitative use of the assay. We have previously extended this approach to measurement of mRNA in a given sample, a technique referred to throughout as qRT-LAMP (Remmel *et al*., 2022; see Figure 1).

**Figure 1:**
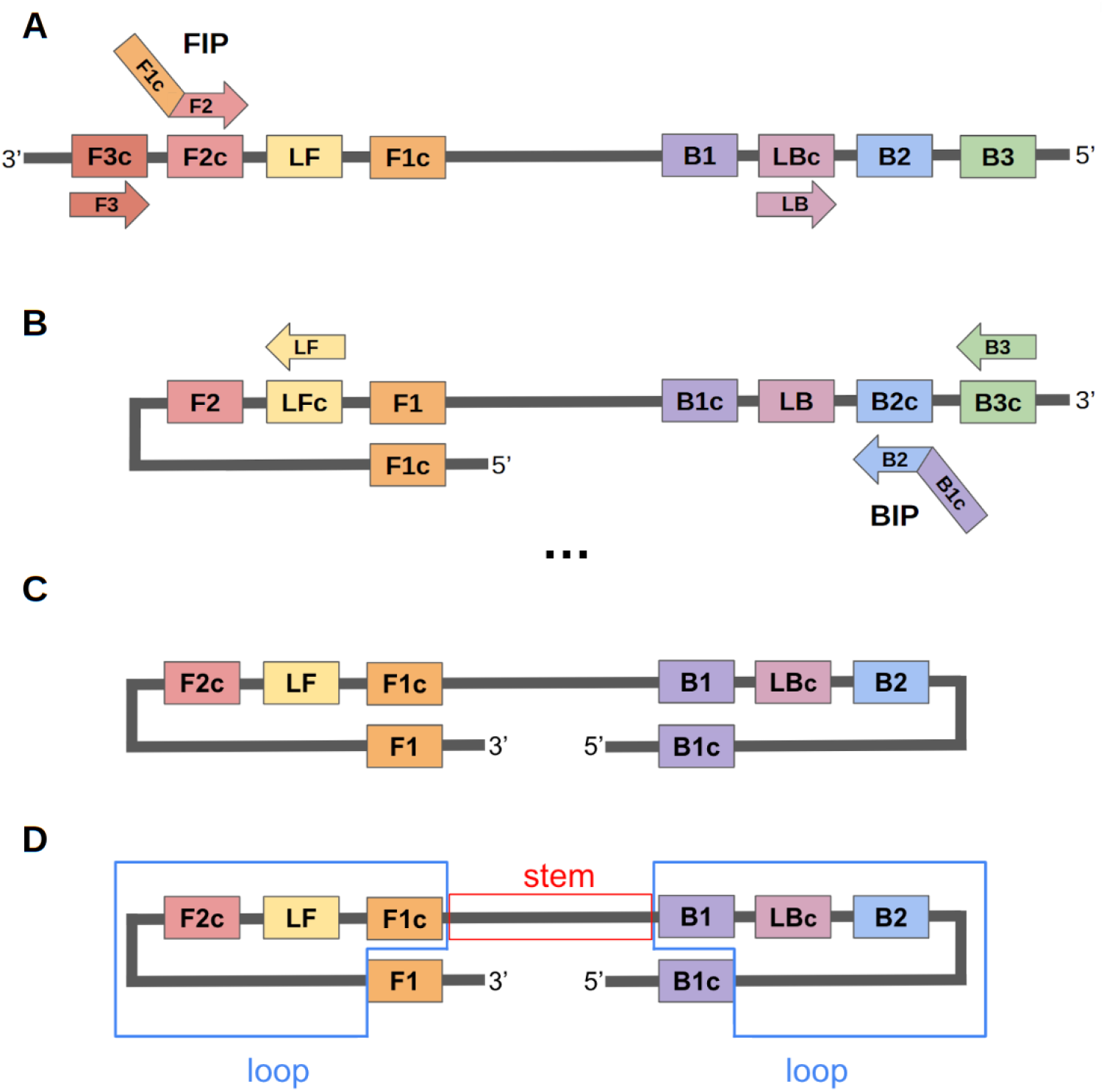
Intermediate steps in the qRT-LAMP reaction. **A**: Following reverse transcription, any of the primers F3, FIP (comprising F2 and F1c sequences), and LB may anneal to the cDNA template. Polymerization from the bound FIP primer leads to introduction of the F1c sequence into the product. **B**: Annealing of the F1c sequence to its complement creates a loop. The primers B3, BIP (comprising B2 and B1c sequences), and LF anneal to this single-loop template. **C**: Annealing of the B1c sequence to its complement creates another loop, resulting in the canonical “dumbbell” LAMP template, or amplicon. **D**: The stem region of the dumbbell template (the span between F1c and B1 sequence regions) contains the target sequence of interest while the loop regions (the regions spanning from F1c/B1 (inclusive) to F1/B1c (exclusive)) provide additional sites to initiate polymerization. Crucially, any of the six primers may anneal and initiate polymerization at any point at which a complementary region is available, leading to creation of other products (not shown for simplicity) distinct from the canonical product.

Current understanding of the ideal properties of primers and targets to optimize quantitative reverse-transcription loop-mediated isothermal amplification (qRT-LAMP)’s performance is incomplete and qualitative; primer design relies on a small set of guiding principles (Panno *et al*., 2020), including restrictions on the GC nucleotide content, melting temperature, and presence of GG repeats in primer sequences. In practice, primer design proceeds in trial-and-error fashion, as these guidelines can often be in conflict and are far easier to simultaneously satisfy for some RNA targets than others. Furthermore, there has been little work to assess the impact on performance of sequence-based and thermodynamic properties of the LAMP primers and amplicon.

Statistical modeling frameworks provide one approach to link assay properties and characteristics of amplification curves. Extensive work exists for modeling both PCR amplification curves (Spiess *et al*., 2008; Matz *et al*., 2013; Subramanian and Gomez, 2014; Nguyen *et al*., 2020) and the effect of primer/target properties on PCR assay performance (Mallona *et al*., 2011; Wright *et al*., 2014; Döoring *et al*., 2019). Previous work on primer design for PCR has identified thermodynamic properties of primer/target interactions (Mann *et al*., 2009; Li and Brownley, 2010; Döoring *et al*., 2019) and sequence length (Huang *et al*., 2022) as key quantities associated with assay specificity. Thermodynamic properties (e.g. free energy of primer-target hybridization) have also been associated with the speed of isothermal reactions (Kimura *et al*., 2011). In contrast, the study of LAMP-based quantitation is still under active development, and methods for qRT-LAMP data analysis are lacking.

Gompertz functions (Gompertz, 1825) have proved a useful tool for empirical modeling of growth curves in varied biological settings (Tjørve and Tjørve, 2017). One formulation of the Gompertz curve is parameterized by an “amplitude” (the maximum value reached at asymptote), a “rate”, and an “offset” (the horizontal shift of a curve along its time axis). Gompertz functions are similar to logistic functions, but allow for steeper increases from baseline and slower approaches to asymptote than logistic functions (Figure 2D). Bayesian and hierarchical Bayesian formulations of Gompertz curves have appeared in biomedical applications (Wiper *et al*., 2010; Sasaki and Kondo, 2016; Tutkun and Demirhan, 2016; Gotuzzo *et al*., 2019; Vaghi *et al*., 2020; Berihuete *et al*., 2021) and have started to appear in LAMP reaction modeling for detection (Carvalho *et al*., 2021). However, no Bayesian or hierarchical methods have yet been applied to characterize qRT-LAMP dynamics or to model the effects of different properties of LAMP targets/primers on quantitative assay performance.

**Figure 2:**
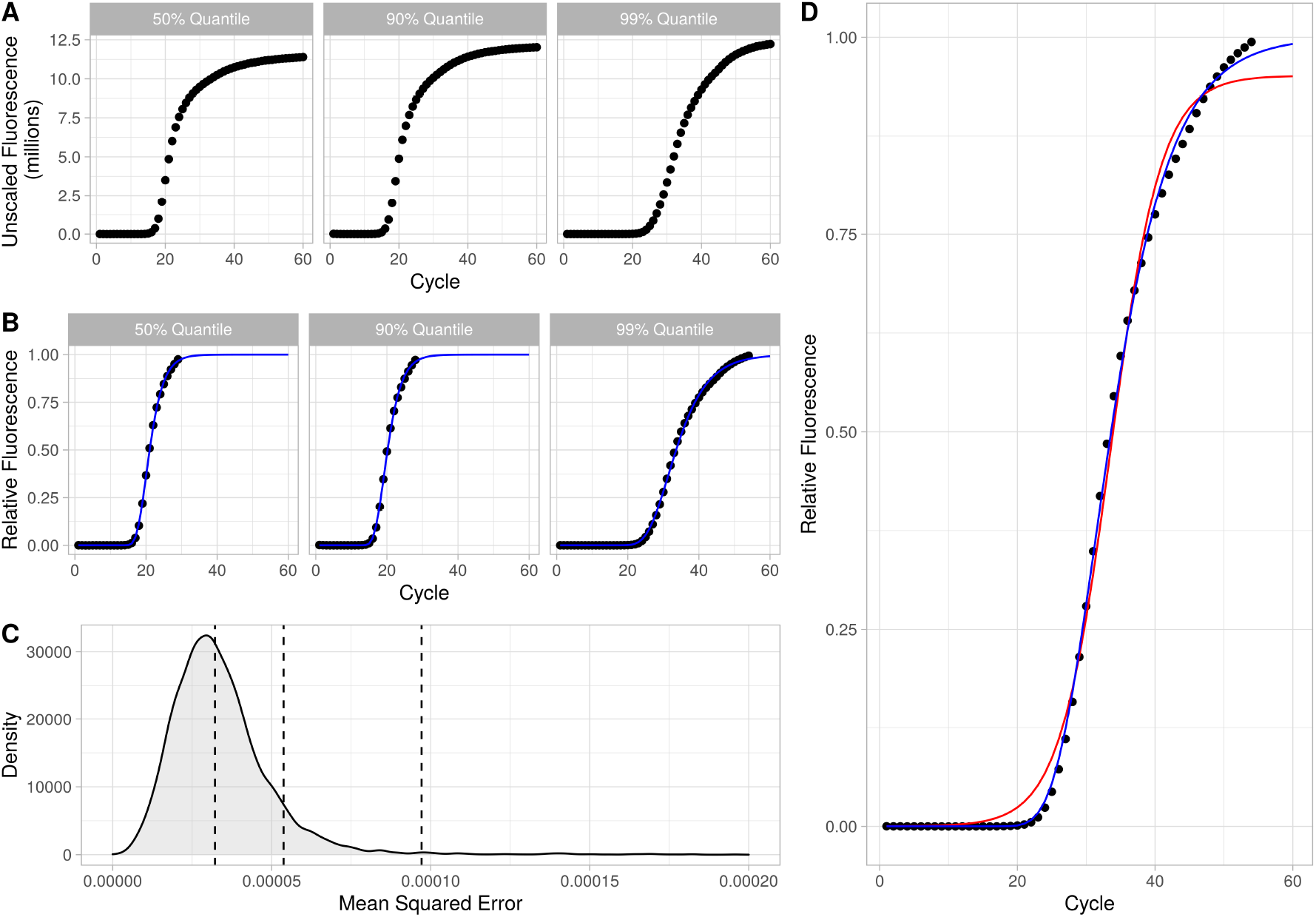
The normalization procedure described in 2.2, with examples from our clinical dataset. **A**: Baseline-corrected LAMP curves with points corresponding to individual fluorescence measurements over 20s pseudo-cycles. **B**: Rescaled and pruned LAMP amplification curves. Points show scaled measurements while solid blue lines correspond to Gompertz curve fits from our iterative procedure. **C**: Distribution of goodness-of-fit (measured by mean squared error) for our fitted Gompertz curves, with 50%, 90% and 99% quantiles indicated by dashed vertical lines. Representative fits from these quantiles are shown in A and B. **D**: Reproduction of the third figure of panel B, with a logistic curve fit for comparison (red).

Here we present a model-based analysis of the quantitative characteristics of the LAMP assay. We first develop a Bayesian modeling framework for inferring the impact of changes in RNA input on amplification rate and offset, and we apply this framework to a dataset of qRT-LAMP curves derived from 80 patients suspected of acute infection. Then, we compile a collection of primer/target features, and extend our hierarchical framework to detect associations between these features and resolution (or the ability to detect differences in RNA input) of the qRT-LAMP assay. We further investigate the effects of primer/target features on assay resolution by generating a novel dataset from synthetic *in vitro*-transcribed (IVT) RNA templates assayed with systematically varied LAMP primer sequences, and by applying our framework to these data. Our model-based analysis highlights several key considerations in the design and optimization of the qRT-LAMP assay.

## 2 Methods

### 2.1 Datasets and technologies

We produced two datasets, one derived from clinical samples and one from synthetic IVT RNA templates. Our clinical dataset comprised remnants of previously-collected PAXgene RNA (Ram-Mohan *et al*., 2022) blood samples collected from 80 patients, each adjudicated to be either bacterially infected (N=21), virally infected (N=22), or non-infected (N=37). Patients samples were taken from two trials (clinicaltrial.gov/NCT03744741: “HostDx Sepsis in the Diagnosis and Prognosis of Emergency Department Patients With Suspected Infections: a Multicenter Pilot Study”, and EudraCT/2020-001039-29: “Efficiency in Management of Organ Dysfunction Associated with Infection by the Novel SARS-CoV-2 Virus Through a Personalized Immunotherapy Approach: The ESCAPE Clinical Trial: The ‘ESCAPE’ Trial (a.k.a ESCAPE)”). At enrollment, patient blood was collected and immediately stored with PAXgene RNA stabilization. For each individual, gene expression was measured for 32 target RNA transcripts: 29 genes previously identified as targets of interest for bacterial/viral/non-infected classification (He *et al*., 2021) as well as 3 housekeeper genes (KPNA6, RREB1 and YWHAB). Two independent aliquots of each sample underwent RNA extraction for profiling by qRT-LAMP and were dispensed along with primers and reagents and run on a 96-well plate with a QuantStudio™ 5 Real-Time PCR System for Human Identification (Applied Biosystems; Waltham, MA USA). We tracked fluorescence for 20 minutes or 60 arbitrary pseudo-cycles (1 cycle = 20 seconds). Another aliquot was used for mRNA quantification using the NanoString (NS) nCounter system. The NS nCounter system directly counts RNA transcript number by measurement of fluorescence via hybridization of capture and reporter probes. As such, higher NS values correspond to higher target expression. We normalized NS measurements by first rescaling counts such that housekeeper genes had a geometric mean of 1000 and then applying a log_2_ transformation to the rescaled counts. To measure mRNA expression with qRT-LAMP, we designed a set of six LAMP primers (FIP/BIP, F3/B3, and LF/LB; Figure 1) for each target using PrimerExplorer v5 (Eiken, 2019) or via a third-party service provider (Primer Digital, Ltd; Helsinki, Finland). qRT-LAMP reactions occurred at 65^*°*^C, in the presence of 90mM KCl and 8mM MgSO_4_. Of the 29 targets of interest, we excluded 4 for technical reasons: FURIN and S100A12 due to the use of different primer concentrations; CTSB and DEFA4 as LF/LB primers did not anneal to their target sites.

Our IVT RNA dataset (preparation described in Remmel *et al*. (2022)) comprised measurements for three targets: IFI27, JUP and OASL. We generated this dataset by assaying different concentrations of synthetic RNA templates corresponding to subsequences of the full-length target sequence of the three different targets. To test the effects of varying different primer properties on qRT-LAMP performance, we varied FIP and BIP primer sequences of each target. These sequence alterations consisted of variations in the lengths of annealing sequence for either the F1c (B1c) or F2 (B2) subsequences of the FIP (BIP) primer (Figure 1). F3, B3, LF, and LB primer sequences were not varied. For each target/primer set combination, we collected qRT-LAMP measurements at three RNA concentrations (10^1^, 10^3^, and 10^5^ copies/*μ*L; 20*μ*L per reaction), with at least two replicates per concentration. We investigated effects of changes in FIP/BIP primer sequence in 35, 48 and 63 primer combinations for IFI27, JUP, and OASL respectively. Reaction conditions for qRT-LAMP with IVT RNA templates were identical to those used for the clinical samples. Replicate samples were dispensed along with primers and reagents and run on a 384-well plate with the QuantStudio™ 5 instrument. For the IVT samples, we tracked fluorescence for 30 minutes or 90 pseudo-cycles.

### 2.2 Amplification curve pre-processing and normalization

To prepare qRT-LAMP curves for analysis, we performed a series of pre-processing steps. We first applied baseline correction to set the initial fluorescence of each reaction approximately to zero. Signal baseline was estimated by identifying a time interval with near-linear signal (2nd differential close to 0), then fitting a line with a Theil-Sen estimator. After correcting the signal baseline, we removed time points with negative fluorescence from each reaction curve.

We then used a nonlinear least-squares approach to fit Gompertz functions to each amplification curve. The Gompertz function is parameterized by three quantities (*a*, *b*, and *c* in Equation 1) associated with the amplitude, rate, and cycle offset, respectively, of the amplification curve. One can think of the *c* parameter as a time-to-threshold measurement (as used for quantitative polymerase chain reaction (qPCR)) in which the threshold is specific to each reaction curve. Higher cycle offset (a right-shifted curve) would indicate lower amounts of RNA, and vice versa. The Gompertz function is defined as follows:

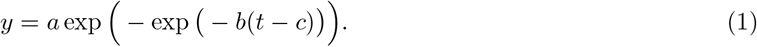

Here, *y* corresponds to baseline-corrected fluorescence and *t* is time in pseudo-cycles. Our procedure estimates 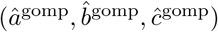 parameters for a given amplification curve. After noting some misfit to the asymptote in early versions of our curve-fitting procedure and motivated by the relative importance (as with qRT-PCR) of the cycle offset (*c*) and amplification rate (*b*) for quantitation, we developed and applied an iterative procedure, described in Algorithm 1, to remove observations exceeding the fitted asymptote at each iteration. We then retained the fitted asymptote 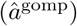 at the last iteration (when no points of the curve exceeded the asymptote) to rescale the curve. A selection of example outputs from this procedure appear in Figure 2.

### 2.3 Single-target model

To investigate the effects of RNA abundance on qRT-LAMP amplification curves for individual targets, we model the Gompertz curve parameter estimates produced by our nonlinear fitting procedure. For samples *i* = 1*,…, N*, replicates *j* = 1*,…, J*, with NS-measured gene expression *x_i_* (a proxy for true RNA input in clinical samples), we model estimated parameters for rate 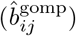 and cycle offset 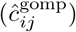 of an individual target as realisations of independent random variables 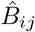 and 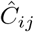 distributed according to:

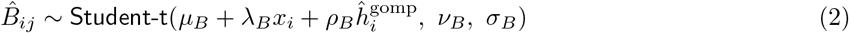

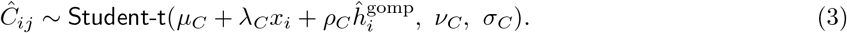

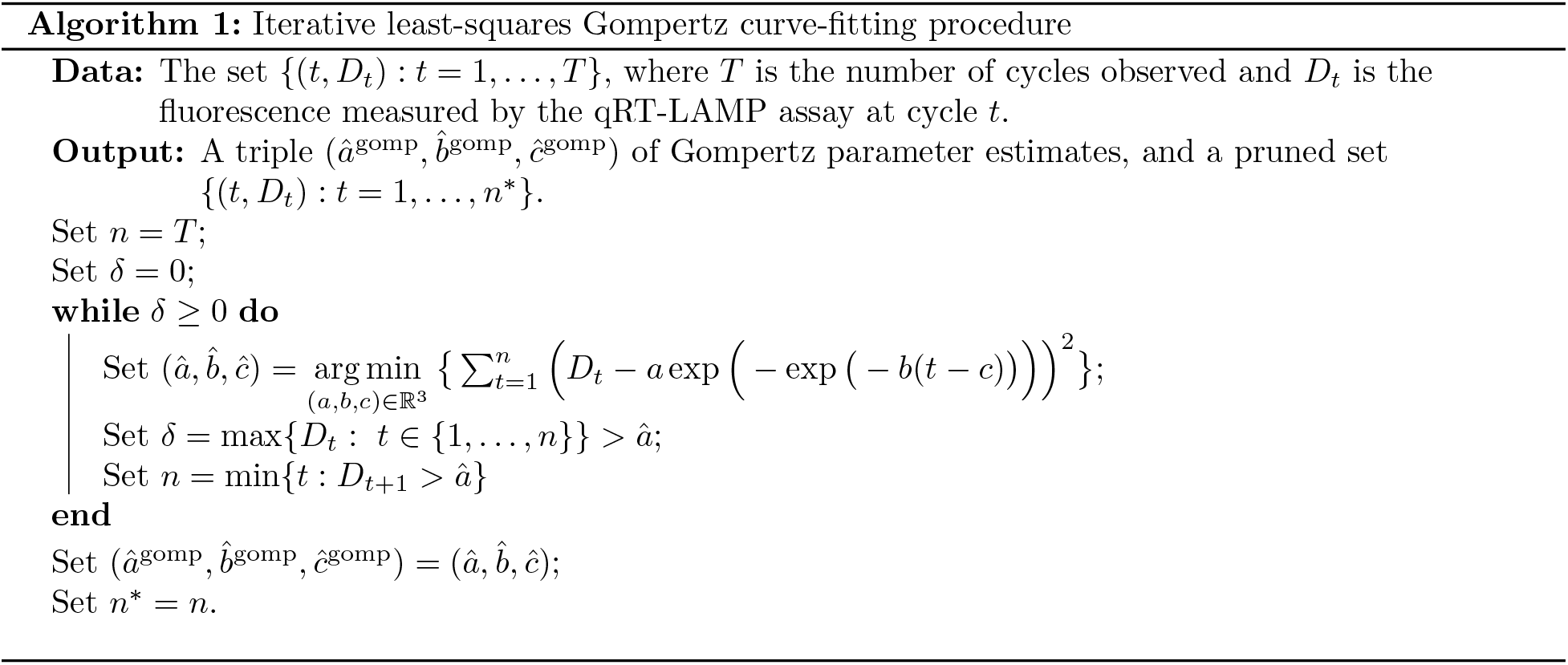

We use a `Student-t`(*μ, ν, σ*) likelihood to provide robustness to outliers, with *μ*, *ν*, and *σ* corresponding to the location, degrees of freedom and scale of the distribution respectively. We refer to *λ_B_* and *λ_C_* as resolution parameters since a larger absolute value of *λ_B_* or *λ_C_* would reflect greater sensitivity, on average, to changes in RNA input for a given target. The input 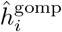 is the geometric mean of the values 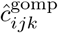 across all replicates *j* and all housekeeper genes *k* ∈ {KPNA6, RREB1, YWHAB}, included to normalize for sample-to-sample variation in RNA yield. The parameters *σ_B_* and *σ_C_* capture variation in rate and cycle offset that is not attributable to variation in RNA input. Bayesian inference is performed with the R package brms (Buörkner, 2017). We specify priors as follows: for the rate model (eq. 2) – *μ_B_* ~ `Norm`(1, 1^2^), *λ_B_* ~ `Norm`(0, 2^2^), *ρ_B_* ~ `Norm`(0, 2^2^), *ν* ~ Γ(2, 0.1), *σ_B_* ~ `Student-t`(3, 0, 2.5^2^); for the cycle offset model (eq. 3), we use matching priors save that *μ_C_* ~ `Norm`(23, 5^2^). We approximate model posteriors with 4000 samples (1000 warm-up) from 3 Hamiltonian Monte Carlo (HMC) chains, assessed for convergence with all parameter posteriors achieving 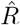 and bulk and tail effective sample sizes above 1000.

### 2.4 Multi-target model

To evaluate associations between primer/target properties and assay resolution, and to model qRT-LAMP amplification curves across all targets, we propose a hierarchical model of the estimated cycle offset. We derive an estimate of the offset 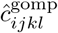 for sample *i*, replicate *j*, target *k*, and (in the case of IVT RNA templates) primer set *l* from our non-linear least squares fitting procedure. We then model the 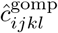 as being independent realisations of random variables 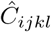, where:

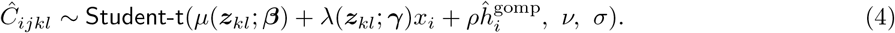

Here, the location of the likelihood comprises RNA-independent baseline offset (*μ*) and RNA-dependent resolution (*λ*) functions, parameterized by ***β*** and ***γ***, respectively. These functions take as an input a vector of sequence-based and thermodynamic features, ***z***_kl_, associated with the given primer set *l* and target *k*. In our clinical dataset, ***z*** has no dependence on *l* as each target corresponds to a single primer set. Also, for our clinical dataset, we let baseline offset *μ* vary across targets *k* by specifying it as a hierarchical random effects term. We also remove dependence of *μ* on ***z***_k_:

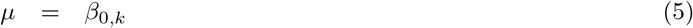

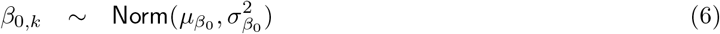

This random term models target-specific effects unrelated to assay characteristics. Our models of baseline offset in clinical samples did not have slopes (*β_z_*) varying by target as we had only one primer set per target.

For our IVT RNA dataset, we note that: *a) x_i_* represents the log RNA copy number at concentration *i*, and *b)* we drop the housekeeper term 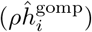 as input RNA concentrations are known. For both our IVT RNA and clinical data, we use *λ* to model effects of assay features on (RNA-dependent) assay resolution (e.g. *λ*(***z***_kl_; *γ*) = *γ*_0_ + *γ_Lstem L,k,l_*).

For analysis of our IVT RNA dataset, we specify two types of models, which we refer to as *across-target* and *within-target* models respectively. For both model types, the RNA-independent offset function, *μ*, includes random intercepts and random slopes to model dependence on assay properties. For example, when using stem length as an input: *μ*(***z***_kl_; ***β***) ≔ *β*_0,k_ +*β*_*L* stem,k_ *L_stem,k,l_*. This is to allow maximum flexibility in the RNA-independent offset to focus on changes in resolution (via *λ*) due to primer set features. The form of the resolution function *λ* for our *across-target* model is identical to that applied to clinical data, while for the *within-target* model, we include random target-specific intercepts in the resolution function *λ*. Again, using stem length as an example, the *within-target* model *λ* function would be:

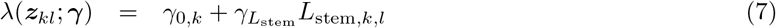

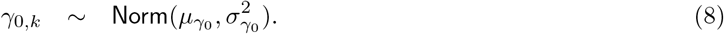

When the *λ* function parameter ***γ*** is a “global” parameter (i.e. not specified as a hierarchical term), we specify independent `Norm`(0, 2^2^) priors for its component parameters. When ***γ*** includes hierarchical terms for the intercept, we specify priors *μ_γ0_* ~ `Norm`(0, 2^2^), *σ_γ0_* ~ `Student-t`(3, 0, 2.5^2^). Priors for the baseline offset parameters ***β*** are set similarly, aside from the intercept term *β*_0_: *μ_β0_* ~ `Norm`(23, 5^2^). The parameters *ν* and *σ* are given priors *ν* ~ Γ(2, 0.1), *σ* ~ `Student-t`(3, 0, 2.5^2^). For multi-target model parameter inference, we use 3 HMC chains of 4000 samples (1000 warm-up), again assessing convergence by confirming that all chains have 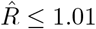 and effective bulk and tail sample sizes above 1000.

### 2.5 Properties of primers and targets

Multiple properties are known to impact effectiveness of LAMP primers, including melting temperature, GC content and GG repeat abundance (Eiken, 2019). In practice, many of these properties are tightly controlled in primer design workflows, resulting in little variability with respect to these quantities. We therefore focused on three classes of LAMP properties: *a)* sequence features of the LAMP target or amplicon; *b)* thermodynamic properties of primers and their targets; and *c)* measures of primer sequence complexity. The full list of properties appears in Table 1. Stem length (*L*_stem_) and loop length (*L*_loop_) measure the total length in nucleotide bases of the amplicon stem and loop regions, respectively (Figure 1D). GC content (*GC*) is the proportion of bases of the entire amplification region (F3 to B3, inclusive) that are either G or C.

**Table 1:** Properties of targets/primers used in hierarchical modelling of LAMP performance.

| Property | Symbol | Unit |
| --- | --- | --- |
| Amplicon stem length | $L_s^{\text{amp}}$ | bp |
| Amplicon loop length | $L_l^{\text{amp}}$ | bp |
| Amplicon GC Content | $GC^{\text{amp}}$ | - |
| 4-complexity (primer $p$) | $C^p$ | - |
| Free energy of annealing (primer $p$) | $\Delta G_a^p$ | $\text{kJmol}^{-1}$ |
| Free energy of self-dimerization (primer $p$) | $\Delta G_{\text{sd}}^p$ | $\text{kJmol}^{-1}$ |
| Free energy of secondary structure formation (primer $p$) | $\Delta G_{\text{ss}}^p$ | $\text{kJmol}^{-1}$ |

Each free energy property measures the strength of some binding process via Gibbs free energy. Free energy of annealing 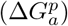 measures the binding affinity of a primer with its complementary target region, while free energy of self-dimerization 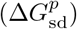 quantifies a primer’s propensity to anneal to another copy of the same primer. Finally, 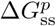 measures a single primer’s tendency to form secondary structure within itself. We follow the approach of Döoring *et al*. (2019), using UNAFold (Markham and Zuker, 2008) for free energy calculations and applying ion concentration adjustments as described in Peyret (2000). For FIP (BIP) primers, we calculated separate free energies of annealing for F1c (B1c) and F2 (B2) primer subsequences.

To measure the extent to which a primer sequence contains unexpected *k*-mer sequences compared to a reference sequence, we also computed sequence complexity (*C^p^*) (Nielsen *et al*., 2003; Xia *et al*., 2010). This measure is based on the information-theoretic notion of entropy (Shannon, 1948). We measure complexity with respect to 4-mers and use the entire human exome as contained in the *Ensembl* database (Yates *et al*., 2020) as our set of reference sequences. We use a simplified formulation of 4-complexity:

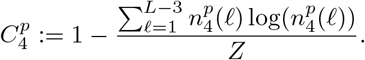

Here, 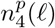 refers to the count of 4-mers found in the reference sequence set that match the 4-mer in primer *p*’s sequence starting at position *f*. The normalising constant *Z* is set to ensure that complexity varies between 0 and 1. A low complexity score indicates that the majority of a primer’s constituent *k*-mers occur commonly in the reference set.

### 2.6 Bayesian stacking for model comparison and validation

To compare multi-target models and identify which LAMP properties are most robustly associated with resolution, we use Bayesian model stacking based on Pareto-smoothed importance sampling approximations of each model’s performance in leave-one-out cross-validation (LOO-PSIS; implemented by the R package loo (Vehtari *et al*., 2020)).

In traditional model stacking, predictions from multiple models are ensembled via a linear combination in order to minimize the average cross-validation prediction error (Wolpert, 1992). The same process is applied in Bayesian model stacking, but rather than using the squared error of models’ predictions, one calculates log predictive densities for left-out observed values (Yao *et al*., 2018). The outputs of stacking are model weights associated with each individual model in the ensemble. These weights will sum to one, and a larger weight indicates a more substantial contribution for the corresponding model (relative to other models in the ensemble) to the optimal model combination. Thus, we take large stacking weights to be indicative of stronger associations of the corresponding model’s features with qRT-LAMP assay resolution. We use Pareto-smoothed importance sampling to approximate (Vehtari *et al*., 2017, 2021) the log posterior predictive density of each left out sample. Pareto *k* diagnostic values > 0.5 would indicate a poor approximation of the LOO posterior predictive density for some observations. We did not observe any such problematic Pareto *k*-estimates in our analyses.

## 3 Results

### 3.1 RNA dependence for qRT-LAMP cycle offset and amplification rate varies widely between targets

Our first aim was to infer the resolution (*λ_B_, λ_C_*) and precision (*σ_B_*, *σ_C_*) parameters associated with RNA input’s effect on qRT-LAMP curve characteristics in clinical samples. We used our single-target models (Equations 2 and 3, Section 2.3) to decompose variation in cycle offset 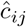; median (across all targets) – 22.2, IQR (across all targets) – 5.03) and rate (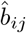; median (across all targets) – 0.334, IQR (across all targets) – 0.076) into separate RNA-dependent and RNA-independent components. We fitted these models for each of our 25 targets of interest. Summaries of the posterior distributions for rate and cycle offset parameters appear in Tables 2 and 3.

**Table 2:** Posterior mean estimates of cycle offset resolution, precision and housekeeper normalization parameters for single-target models based on each target (values in brackets = 95% credible intervals).

| Target | $\lambda_C$ | $\sigma_C$ | $\rho_C$ |
| --- | --- | --- | --- |
| ARG1 | -0.77 (-0.84, -0.71) | 0.61 (0.50, 0.74) | 1.18 (1.05, 1.30) |
| BATF | -1.14 (-1.31, -0.97) | 0.68 (0.59, 0.77) | 0.88 (0.78, 0.99) |
| C3AR1 | -0.87 (-0.97, -0.77) | 0.56 (0.46, 0.68) | 0.59 (0.49, 0.69) |
| C9orf95 | -0.74 (-0.99, -0.49) | 0.85 (0.74, 0.98) | 0.55 (0.42, 0.68) |
| CD163 | -0.62 (-0.70, -0.55) | 0.43 (0.35, 0.50) | 0.40 (0.33, 0.47) |
| CEACAM1 | -0.68 (-0.78, -0.59) | 0.88 (0.74, 1.01) | 0.54 (0.40, 0.68) |
| CTSL1 | -1.94 (-2.16, -1.71) | 1.30 (1.07, 1.53) | 1.14 (0.95, 1.34) |
| GADD45A | -0.75 (-0.90, -0.61) | 0.96 (0.75, 1.14) | 0.32 (0.17, 0.47) |
| GNA15 | -0.82 (-1.02, -0.62) | 0.62 (0.54, 0.71) | 0.97 (0.87, 1.06) |
| HK3 | -0.74 (-0.81, -0.66) | 0.41 (0.36, 0.46) | 0.72 (0.66, 0.78) |
| HLA-DMB | -0.81 (-1.36, -0.25) | 3.13 (2.79, 3.53) | 0.73 (0.37, 1.08) |
| IFI27 | -0.94 (-1.05, -0.83) | 1.80 (1.51, 2.12) | 0.66 (0.42, 0.90) |
| ISG15 | -0.82 (-0.95, -0.68) | 1.62 (1.37, 1.90) | 0.91 (0.69, 1.14) |
| JUP | -1.49 (-1.73, -1.24) | 2.36 (2.04, 2.69) | 1.50 (1.20, 1.81) |
| KCNJ2 | -0.77 (-0.90, -0.65) | 0.52 (0.41, 0.66) | 0.14 (0.05, 0.24) |
| KIAA1370 | -0.25 (-0.50, -0.00) | 0.74 (0.60, 0.89) | 0.45 (0.32, 0.57) |
| LY86 | -0.82 (-0.98, -0.66) | 0.55 (0.45, 0.64) | 0.88 (0.80, 0.97) |
| OASL | -0.89 (-0.96, -0.81) | 0.65 (0.53, 0.78) | 0.87 (0.75, 0.99) |
| OLFM4 | -1.07 (-1.19, -0.95) | 1.50 (1.31, 1.71) | 0.56 (0.35, 0.78) |
| PDE4B | -0.90 (-1.26, -0.53) | 1.32 (1.17, 1.49) | 0.99 (0.80, 1.18) |
| PER1 | -1.41 (-1.78, -1.05) | 2.07 (1.72, 2.45) | 0.61 (0.32, 0.89) |
| PSMB9 | -0.88 (-1.02, -0.74) | 0.51 (0.41, 0.62) | 0.86 (0.77, 0.94) |
| RAPGEF1 | -0.88 (-1.36, -0.40) | 1.26 (1.02, 1.51) | 0.97 (0.75, 1.17) |
| TGFB1 | -0.66 (-0.75, -0.58) | 0.53 (0.45, 0.62) | 0.85 (0.76, 0.94) |
| ZDHHC19 | -2.58 (-2.95, -2.25) | 2.85 (2.10, 3.70) | 1.32 (0.96, 1.69) |

**Table 3:** Posterior mean estimates of rate resolution, precision and housekeeper normalization parameters for single-target models based on each target (values in brackets = 95% credible intervals).

| Target | $\lambda_B$ | $\sigma_B$ | $\rho_B$ |
| --- | --- | --- | --- |
| ARG1 | 0.02 (0.02, 0.02) | 0.03 (0.02, 0.03) | -0.03 (-0.03, -0.02) |
| BATF | 0.03 (0.02, 0.03) | 0.02 (0.02, 0.03) | -0.02 (-0.02, -0.01) |
| C3AR1 | 0.02 (0.02, 0.02) | 0.02 (0.02, 0.02) | -0.01 (-0.02, -0.01) |
| C9orf95 | 0.02 (0.01, 0.03) | 0.02 (0.02, 0.03) | -0.01 (-0.02, -0.01) |
| CD163 | 0.00 (0.00, 0.01) | 0.02 (0.02, 0.02) | -0.01 (-0.01, -0.01) |
| CEACAM1 | 0.00 (0.00, 0.01) | 0.02 (0.02, 0.02) | -0.01 (-0.02, -0.01) |
| CTSL1 | 0.03 (0.02, 0.03) | 0.03 (0.02, 0.03) | -0.01 (-0.02, -0.01) |
| GADD45A | 0.02 (0.01, 0.02) | 0.02 (0.02, 0.03) | -0.02 (-0.02, -0.01) |
| GNA15 | 0.01 (0.00, 0.02) | 0.02 (0.01, 0.02) | -0.02 (-0.02, -0.01) |
| HK3 | 0.00 (0.00, 0.00) | 0.02 (0.02, 0.02) | -0.01 (-0.02, -0.01) |
| HLA-DMB | 0.01 (0.00, 0.01) | 0.03 (0.03, 0.03) | -0.01 (-0.01, -0.01) |
| IFI27 | 0.01 (0.00, 0.01) | 0.02 (0.02, 0.02) | -0.01 (-0.01, 0.00) |
| ISG15 | 0.00 (0.00, 0.01) | 0.02 (0.02, 0.02) | -0.01 (-0.02, -0.01) |
| JUP | 0.02 (0.02, 0.02) | 0.03 (0.02, 0.03) | -0.02 (-0.02, -0.01) |
| KCNJ2 | 0.00 (0.00, 0.00) | 0.01 (0.01, 0.02) | -0.01 (-0.01, 0.00) |
| KIAA1370 | 0.01 (0.00, 0.02) | 0.03 (0.03, 0.04) | -0.01 (-0.02, -0.01) |
| LY86 | 0.02 (0.01, 0.02) | 0.02 (0.01, 0.02) | -0.02 (-0.02, -0.01) |
| OASL | 0.01 (0.01, 0.02) | 0.02 (0.02, 0.02) | -0.02 (-0.02, -0.01) |
| OLFM4 | 0.01 (0.01, 0.02) | 0.03 (0.03, 0.04) | -0.01 (-0.01, 0.00) |
| PDE4B | 0.01 (0.00, 0.02) | 0.02 (0.02, 0.02) | -0.01 (-0.02, -0.01) |
| PER1 | 0.04 (0.03, 0.05) | 0.05 (0.04, 0.06) | -0.01 (-0.02, -0.01) |
| PSMB9 | 0.01 (0.00, 0.01) | 0.01 (0.01, 0.02) | -0.01 (-0.02, -0.01) |
| RAPGEF1 | 0.00 (0.00, 0.01) | 0.02 (0.02, 0.02) | -0.02 (-0.02, -0.01) |
| TGFB1 | 0.01 (0.00, 0.01) | 0.02 (0.01, 0.02) | -0.01 (-0.02, -0.01) |
| ZDHHC19 | 0.04 (0.04, 0.05) | 0.04 (0.04, 0.05) | -0.02 (-0.02, -0.01) |

We see a wide range of inferred posterior means for both *λ_C_* and *σ_C_*. For the parameter *λ_C_*, which denotes resolution with respect to cycle offset, larger negative values correspond to improved resolution. Offset precision *σ_C_* is always positive, with lower values corresponding to a more precise assay. For example, targets CTSL1 and ZDHHC19 both show good resolution (posterior means of −1.94 and −2.58, 95% credible intervals of (−2.16, −1.71) and (−2.95, −2.25), respectively), but different levels of precision (posterior means of 1.30 and 2.85, 95% credible intervals of (1.07, 1.53) and (2.10, 3.70), respectively). Inferences of poor resolution and precision for some targets (e.g. KIAA1370) indicate the potential need for further optimization of assay properties and/or conditions. We also note that, as expected and as reflected by the posteriors for *ρ_C_*, the geometric mean of the housekeeper cycle offsets is generally correlated with the cycle offset of the target (i.e., lower sample yield should correspond to higher cycle offsets for all targets).

The amplification rate parameter *λ_B_* also varied between targets, with a larger positive value indicating greater average sensitivity to changes in RNA input. While many reactions, including for example ISG15 and HK3, showed almost no detectable variation in rate due to RNA input (posterior mean for *λ_B_* of 0.00, 95% credible intervals of (0.00, 0.00) in both cases), others such as PER1 and ZDHHC19 did show a dependence on NS-derived measurements of RNA (posterior means for *λ_B_* of 0.04 in both cases, 95% credible intervals of (0.03, 0.05) and (0.04, 0.05) respectively). Posterior summaries for the precision in amplification rate (*σ_B_*) were roughly similar across targets. We also found that housekeeper levels showed a negative association with the rate of amplification (for example, 95% posterior credible intervals for *ρ_B_* of (−0.03, −0.02) in ARG1 and (−0.02, −0.01) in BATF).

To further elucidate the range of behaviours captured by these single-target models, we simulated from the posterior predictive distributions of 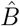 and 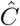 for four targets (CEACAM1, BATF, PER1, and ZDHHC19), to parameterize scaled Gompertz curves. We used these samples to simulate scaled Gompertz curves from targets showing different types of association between RNA input and amplification rate/cycle offset. We simulated these curves at three NanoString-derived RNA input values (results shown in Figure 3). CEACAM1 displays negligible rate dependence on RNA input, whereas all three others appear qualitatively different in shape across the gene expression range. *PER1* and *ZDHHC19* exhibit the greatest offset resolution, but this is accompanied by substantially worse precision. A key point to emphasize is that, unlike in the case of qPCR, opting for an approach based purely on time-to-threshold will under-utilize the full information content of qRT-LAMP curves. As rates of amplification also appear important for characterizing qRT-LAMP reactions, parameter estimates of the rates could be used in tandem with cycle offsets for target quantitation.

**Figure 3:**
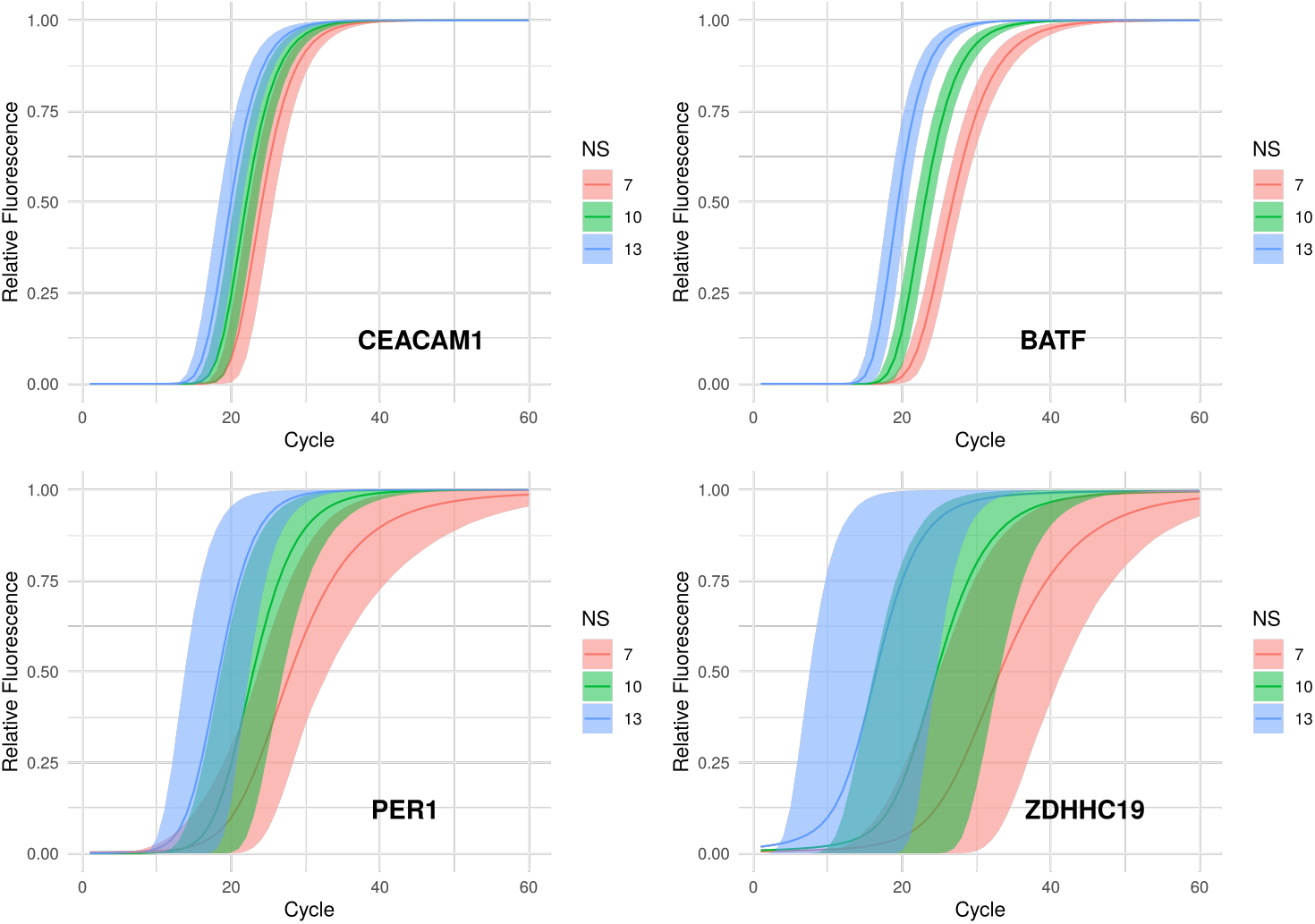
Simulated amplification curves for four primers (CEACAM1, RAPGEF1, BATF, and ZDHHC19). Posterior predictive samples of Gompertz curves of scaled fluorescence based on posterior predictive samples from models 2 and 3. For each target, curves are simulated at three different RNA input (NS) levels – note that higher NS values correspond to higher expression and, generally, to lower cycle offsets (left-shifted amplification curves). Solid lines depict posterior means, while highlighted regions correspond to 95% credible intervals.

### 3.2 Multi-target model identifies assay properties associated with LAMP resolution in patient data

We fitted the multi-target model described in Equation 4 to our clinical dataset using different combinations of primer/target features in the resolution function *λ*(***z***_k_; ***γ***). We considered the following forms for *λ* in our analysis: *a)* an intercept-only model *λ*(***γ***) = *γ*_0_; *b)* for each of the three amplicon properties, a univariate linear function (e.g. for stem length *L*_stem_, *λ*(*z_k_*; ***γ***) = *γ*_0_ + *γ*_*L* stem_ *L*_stem,k_); and *c)* for each primer-specific quantity and primer pair, a bivariate model (e.g. in the case of free energy of annealing for the F1/B1 pair, 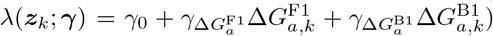 Note again that in this setting we only have one candidate primer set for each target and so no dependence on primer set *l*.

Posterior summaries of the resolution function coefficients as well as the leave-one-out Pareto smoothed importance sampling (LOO-PSIS) stacking weights for each of these models appear in Figure 4. For the stacking weights (left-hand panel of Figure 4), the higher the stacking weight, the higher quality predictions the model was judged to have made for a left-out sample in comparison to alternative models. In the right-hand panel of Figure 4, we show posterior summaries of effect sizes (i.e. ***γ*** coefficients) associated with each of the primer/target properties used as features in the resolution function, *λ*.

**Figure 4:**
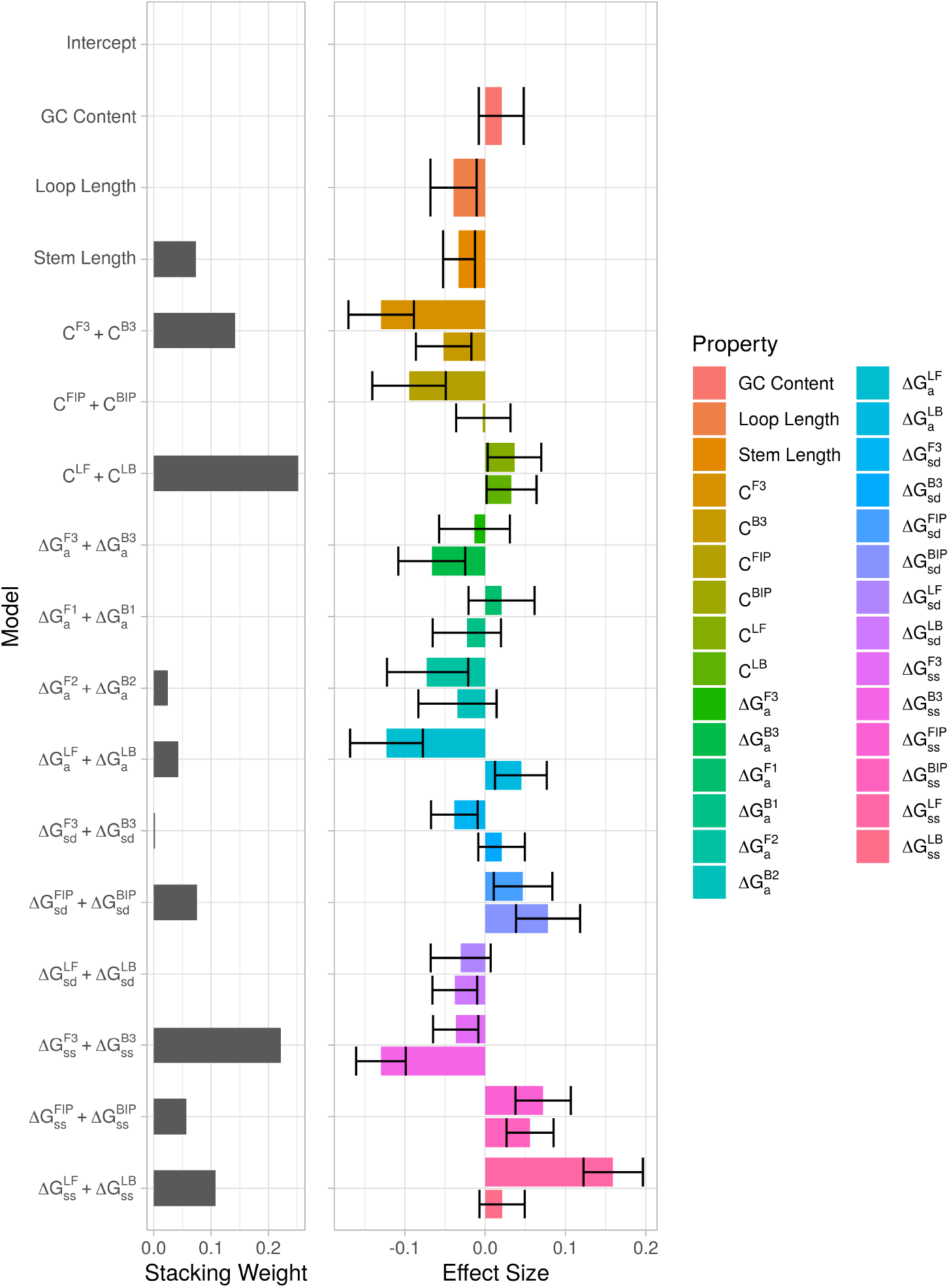
Posterior summaries of multi-target models specified in Sections 2.4 and 3.2, based on combinations of primer/target properties, with their associated Bayesian model stacking weights (left) and ***γ***-coefficient effects sizes (right). Effect sizes are posterior means, with error bars delineating 95% confidence intervals. Notation as given in Table 1.

We found quantities associated with resolution in both positive and negative directions whose corresponding models were assigned strong weightings by the LOO-PSIS procedure. Increased stem length, complexity in the F3/B3 primer pair, free energy of annealing in F2/B2 regions of the FIP/BIP primers, and free energy of secondary structure formation in F3/B3 were associated with improved resolution (negative effect sizes), and all corresponding models received relatively high stacking weights. These associations, particularly for F3, may reflect the importance of the primer to the synthesis of a full-length template, resulting in more sites for primer annealing and amplification (Figure 1). Increases in complexity of LF/LB primers, free energy of self-dimerization for FIP/BIP, and free energy of secondary structure formation in LF/LB were associated with worse resolution (positive effect sizes), with corresponding models also receiving high stacking weights. We do observe some primer pairs for which posterior mean effect sizes were in opposing directions. For example, free energy of self-dimerization shows a positive effect for the B3 primer and a negative effect for F3. We see that no such models were assigned large stacking weights. We might expect these different associations with resolution due to stochasticity of the LAMP assay in the production of intermediates from either strand of the target.

### 3.3 Multi-target model identifies distinct across-target and within-target effects of primer variation

In the previous analysis, we pooled our data across targets to estimate associations of assay properties with resolution. However, while we showed associations of these properties with resolution across targets, we could not determine for any *individual* target whether variation in these properties was associated with variation in resolution. To address the latter objective (arguably more important for LAMP optimization), we generated a IVT RNA dataset comprising multiple primer sets (containing multiple combinations of the FIP and BIP primers) for each target. We fitted models with two different forms of resolution function. In the first form, for each combination of assay properties we fitted a model based on those properties alone with no target-level grouping terms (*across-target* models). In this way, the resolution function *λ* did not depend on target (*k*) other than through the primer set (*l*) features ***z***_kl_ (e.g., *λ*(*z_kl_*; ***γ***) = *γ*_0_ + *γ*_*L* stem_ *L*_stem*,k,l*_). In the second model form (*within-target*), we specified a hierarchical random effects term (varying by target) for the resolution intercept, *γ*_0_. In this way, we were able to infer the dependence of resolution on primer set properties while accounting for intrinsic target-to-target variation in resolution. Summarized inferences and model stacking weights after fitting of the across-target and within-target models to the IVT RNA data appear in Figure 5.

**Figure 5:**
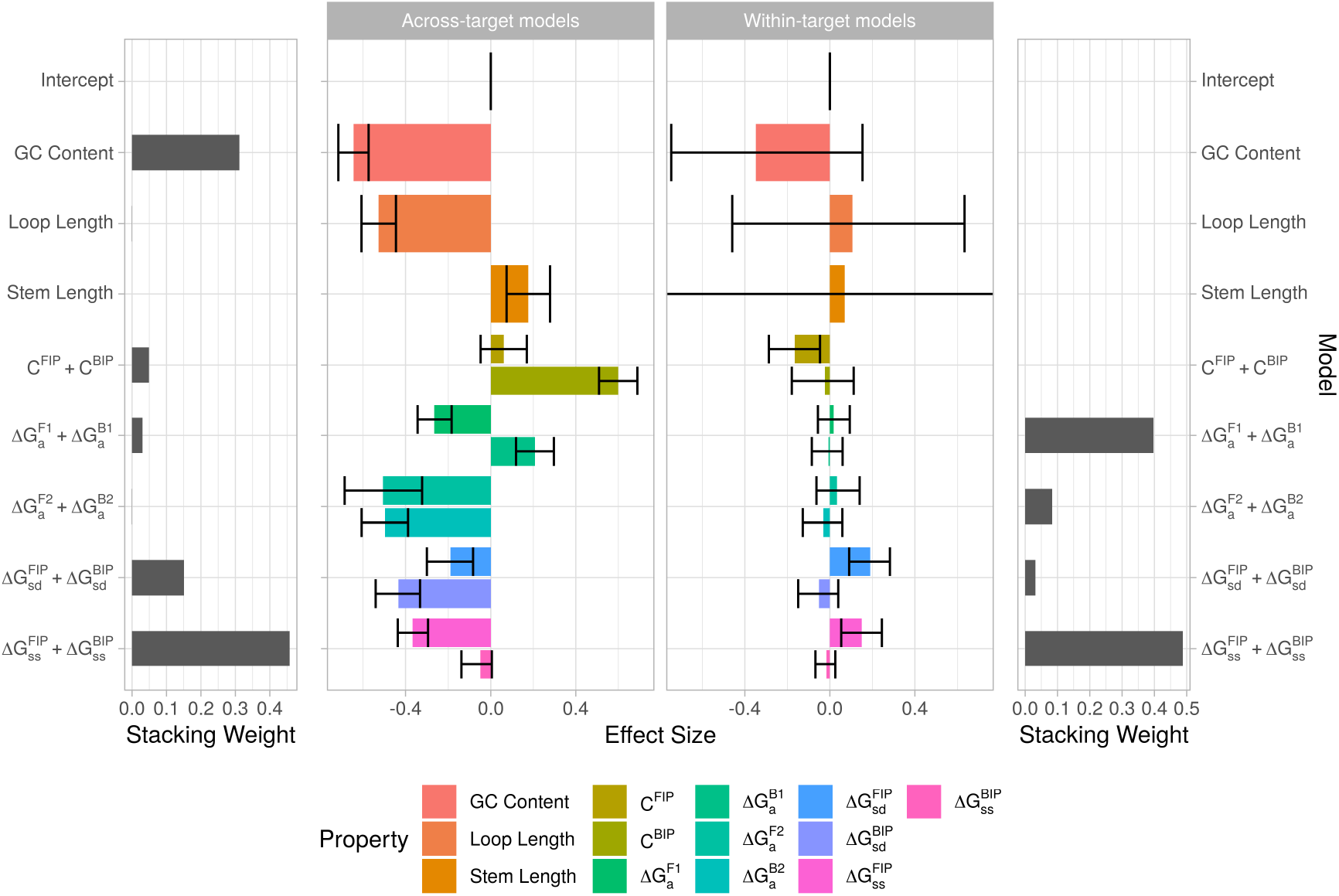
Posterior summaries of multi-target models as specified in Sections 2.4 and 3.3 applied to IVT RNA data. Effect sizes for ***γ***-coefficients from across-target (center left) and within-target (center right) models, as well as associated model stacking weights (far left/right respectively). Error bars delineate 95% confidence intervals.

We note that the across-target models gave far more inferences than the within-target models of high-confidence, high-magnitude effect sizes of assay properties on resolution. Relatively few properties were inferred to have effect sizes of high magnitude (many 95% credible intervals contain 0) for within-target models. Importantly, these results indicate that much of the variation in resolution explained in previous models may be due to intrinsic target-to-target variation not attributable to the assay properties we considered. Of the within-target models, those based on complexity of the FIP primer, free energy of self-dimerization of the FIP primer, and free energy of secondary structure formation of the FIP primer had significant posterior mean effect sizes (i.e. 95% posterior credible intervals not containing zero). Notably, several of these within-target inferences differ in directionality from their across-target counterparts, indicating that not accounting for intrinsic target-to-target variation could be masking the nature of the association between assay properties and resolution. Furthermore, these results highlight the importance of utilizing multiple primer sets per target in characterizing this variation. Interestingly, we do note that both within- and across-target models agreed in allocation of the largest stacking weight to free energy of secondary structure formation for FIP/BIP, a novel quantity for consideration in the design of qRT-LAMP assays.

## 4 Discussion

We have presented and applied a novel model-based framework to characterize quantitative performance of the qRT-LAMP assay. We first introduced a non-linear least-squares fitting procedure based on Gompertz curves that allows for fast, accurate characterization of qRT-LAMP reactions. Based on LAMP reaction curve characteristics derived from this Gompertz curve-based fitting procedure, our Bayesian hierarchical models have generated new insights into the qRT-LAMP reaction and its properties at the single- and multi-target levels. At the single-target level, our analysis identified RNA-independent and RNA-dependent associations with qualitative characteristics and quantitative performance of qRT-LAMP in a dataset of patients with suspected acute infection. Our investigation of the effects of various sequence-based and thermodynamic primer/target properties on qRT-LAMP performance identified novel associations between some of these properties and LAMP assay resolution. We’ve not only adapted our framework to single-target and multi-target settings but also to different qRT-LAMP data modalities (from IVT RNA and clinical samples). Finally, we have used our IVT RNA data and modeling framework to investigate the extent to which associations observed between primer/target properties and LAMP performance will be borne out by artificially altering primers as part of an assay optimization workflow.

We highlight several advantages of our approach. The single-target models introduced in Section 2.3 and applied in Section 3.1 provide a formal framework for inference with uncertainty of qRT-LAMP performance measures such as resolution and precision. This approach allows assay developers to ascertain the quantitative performance of an already-designed LAMP target. Such inferences may in turn motivate optimization of the given target or, in light of poor target performance, alternative target selection as part of a broader panel of target biomarkers. Furthermore, both our single-target and multi-target models allow decomposition of variation in LAMP reaction characteristics into components useful for mRNA quantitation (i.e. amplification rate and cycle offset). By incorporating different features of the assay into model components for RNA-dependent and RNA-independent variation, we can uncover potentially tuneable determinants of LAMP assay performance. Indeed, our framework applied to IVT RNA data revealed significant associations between primer and target features and assay resolution. While IVT RNA data are useful for assay development due to their cost and ease of handling, they are not biologically representative and would likely not recapitulate assay performance seen on clinical samples. As we could adapt our models to both IVT RNA and clinical samples, we were able to compare our inferences across different data types and to better anticipate clinical performance of a given assay. We see our framework and analysis as important first steps towards a system for LAMP assay design and optimization.

However, the results of our hierarchical modeling (Section 3.3) highlight the complexities of optimizing qRT-LAMP assay performance. While FIP and BIP primers are known to play a central role in LAMP-based amplification, our model-based analysis (Figure 5) provides evidence that other primer characteristics may also affect assay performance. Models based on loop amplification and inner primer pairs were identified by model stacking as producing some of the most robust and predictive associations. Furthermore, our findings suggest that increasing a given property (for example, free energy of self-dimerization) may be associated with improved resolution for one primer pair (FIP/BIP) but the opposite for another (LF/LB). As changing one oligo property (e.g. secondary structure formation) may alter another (propensity to self-dimerize), our results support joint consideration of all primer pair characteristics as part of assay optimization efforts. We also observe, for several properties, a difference in directionality of effect sizes between forward and backward primers (Figure 5). Subject to further investigation, we hypothesize that this difference in directionality between members of a primer pair could arise due to the orientation of initial RNA inputs to the LAMP assay. For example, B3 primers could anneal to RNA inputs before reverse transcription while F3 primers – relying on the generation of an F3c region – could not anneal until after reverse transcription, and this could explain some degree of within-primer pair disparity.

While we see advantages to our approach in characterizing pre-designed LAMP assay performance, further work is required to accurately predict performance in a proposed assay for a potentially novel target. As part of our efforts towards a more predictive model, we analyzed multiple primer sets per target with our IVT RNA dataset. Indeed, we found for multiple models that the effects of varying properties of primers and amplicons differ in their strength and direction when viewed across-target vs. within-target (Figure 5). Put another way, accounting for target-to-target differences in resolution not due to the assay features attenuated the strength of association between the features and resolution. These results suggest that other assay features may be explaining target-to-target differences in resolution and that predictive models based on pooling all targets together (i.e. across-target models) may not generalize well to new targets. Interestingly, recent work on predictive modeling of PCR amplification from primer/target features showed good performance in held-out test sets (Döoring *et al*., 2019). This performance could be due to both a focus on a highly related set of targets which might be expected to share assay performance properties as well as due to random data subsampling to create training, validation and test data splits for model development. In this setting, all splits would likely include observations for a given target (a kind of information leakage), complicating interpretation of the model’s generalizability. However, if we consider a highly related family of targets to be similar to a single target, the results of Döoring *et al*. (2019) are consistent with our findings that within-target trends (rather than general trends shared across targets) in the relationship between LAMP assay performance and primer properties are important for optimization of a given target or target family. Reliably determining the optimal properties of a primer set to achieve a pre-specified level of quantitative performance would allow for less trial-and-error in the development process, and ongoing and future work will be required to add both more targets and primer sets to our IVT RNA dataset.

Our work is not without limitations. To extend and more comprehensively validate the results presented here, a follow-up study will be required to profile many more targets with – crucially – multiple combinations of primers for each target. In particular, we note that while our IVT RNA data analysis included multiple primer sets per target, we were only able to assay three targets due to time/cost constraints. Further data collection will also permit investigation of additional candidate assay properties, including gDNA interference, sensitivity to reagents and other sequence-based/thermodynamic quantities. In addition, while IVT RNA data are useful for assay development due to their cost and ease of handling, they are not biologically representative. The degree to which conclusions drawn from IVT RNA translate to clinical samples is the subject of ongoing and future investigation. Also, while in our multi-target modeling work we have investigated variation in resolution with respect to cycle offset, we have not provided a complementary analysis for the other properties shown to be important in our single-target modeling section. These include resolution with respect to amplification rate, and precision. Extending our analysis to identify associations between these performance characteristics and assay properties is the subject of ongoing and future work. In addition, for simplicity, we assumed a linear relationship between assay properties and resolution. Further investigations will involve nonlinear relationships between these quantities.

## 5 Conclusion

This work provides an extensible platform to address the significant knowledge gap in enabling reliable quantitative design and use of the qRT-LAMP assay. Our analysis has uncovered important insights of RNA-dependent and RNA-independent effects on qRT-LAMP assay behavior as well as assay properties associated with improved quantitative resolution of LAMP. Our efforts have also raised important considerations for constructing truly predictive systems of assay performance. With further data collection, expansion of our set of assay features, and extension of our modeling framework, we hope to take another important step towards principled design of robust, clinically performant qRT-LAMP assays.

## Author Contributions

JB and MM developed the modeling framework. MC and DB collected IVT RNA data. JB and MM wrote the paper.

## Acknowledgements

We thank Melissa Remmel, Sabrina Coyle and Yuan Yuan for providing the clinical sample data. We also gratefully acknowledge valuable discussions with Ljubomir Buturovic, Timothy Cannings, Isabella Deutsch, Isabel Johnson, Sara Masarone, Uros Midic, Morton, Amitesh Pratap, Arthur Radley, Torben Sell, Vera Silva, and Tim Sweeney.

